# Cardiac Effects of a Novel pH-Insensitive Sodium-Calcium Exchanger Mouse

**DOI:** 10.1101/2023.05.11.540465

**Authors:** Rui Zhang, Xiaokang Wu, Brian Kim, Catherine Xie, Devina Gonzalez, Raven Norris, Nicholas Chin, Liang Li, Scott John, Kenneth D. Philipson, Michela Ottolia, Joshua I. Goldhaber

## Abstract

**BACKGROUND:** Cardiac sodium-calcium exchange (NCX1) is the dominant calcium (Ca) efflux mechanism in cardiomyocytes and is strongly regulated by pH. However, the role of NCX1 pH sensitivity in normal cardiac function is unknown.

**METHODS:** We used CRISPR/Cas9 to produce a pH-resistant NCX1 mouse by replacing the histidine at position 165 of NCX1 with an alanine (H165A). Hearts were studied using echocardiography and ECG. RNA and protein expression levels were assessed using qPCR and Western blotting. Isolated ventricular cardiomyocytes were loaded with Ca indicators and patch clamped to record intracellular Ca transients and membrane current and voltage.

**RESULTS:** H165A mice live into adulthood with slightly reduced LV systolic function, normal heart rate and shortened QT interval. Both male and female animals exhibit reduced growth, but females eventually reach normal body weight. In patch clamped myocytes, NCX current (I_NCX_) evoked by voltage ramps was reduced by 35% (at +80 mV). Lowering pH_i_ to 6.5 using Na-Acetate had *no effect* on I_NCX_ in H165A myocytes, whereas the same intervention in wildtype (WT) inhibited I_NCX_ by 69% (at +80 mV, p<0.01). There was no change in H165A ventricular cardiomyocyte Ca transients measured with fura-2 AM. However, action potential duration was reduced 68%, consistent with the shorter QT interval. This coincided with a 37% reduction in L-type Ca current and increased expression of repolarizing K^+^ channels. H165A mice are also resistant to ischemia/reperfusion injury.

**CONCLUSIONS:** The H165A mutation attenuates pH regulation of NCX1 in mice, is associated with reduced growth and accelerates cardiac repolarization without compromising excitation-contraction coupling. The mutation also confers cardioprotection. The H165A mouse is the first evidence that pH regulation of NCX1 affects cardiac physiology and is a potential model for studying the role of NCX1 pH-sensitivity on both physiological and pathophysiological cardiac function.

## Introduction

The cardiac sodium-calcium exchanger (NCX1) is the dominant calcium (Ca) efflux mechanism in cardiac cells, removing in diastole precisely the amount of Ca that enters through L-type Ca channels during systole.^1^ The direction and extent of ion transport by NCX1 is constrained by membrane voltage and the relative concentrations of sodium (Na) and Ca in the intracellular and extracellular space. Modest increases in intracellular Na produced by the Na/K ATPase inhibitor digoxin reduce net Ca efflux rate and increase sarcoplasmic reticulum (SR) Ca load and contractility.^2^ Much higher intracellular Na concentrations, particularly when resting membrane potential is depolarized, cause NCX1 to function in reverse, resulting in Ca entry while excess Na is extruded. Thus, cellular regulation of Na at physiological levels is critical for NCX1 to fulfill its Ca efflux role. Indeed, global knockout of NCX1 is embryonic lethal.^3^ Nevertheless, mouse models of NCX1 overexpression and cardiac-specific knockout of NCX1 are viable, demonstrating an impressive capacity for adaptation and compensation by alternative Ca regulatory mechanisms.^4^ For example, cardiac-specific NCX1 knockout mice (constitutive or inducible) are fertile with systolic left ventricular (LV) function in the normal range.^5, 6^ These animals adapt to the reduction in Ca efflux by restricting Ca influx, and increasing Ca efflux through alternative pathways including the plasma membrane Ca ATPase pump (PMCA). NCX1 KO mice are also resistant to arrhythmia and ischemia/reperfusion injury, presumably because they prevent Ca influx and overload during reperfusion.^6, 7^.

NCX1 is strongly regulated by intracellular pH.^8–11^ Small decreases in pH inhibit NCX1 activity.^8–10, 12^ This pH dependence has been implicated in the pathogenesis of ischemia/reperfusion (I/R) injury.^13–16^ However it is unknown if pH regulation of NCX1 affects normal cardiac physiology and function. We reported that substitution of a single histidine to alanine at position 165 (H165A) of the NCX1 protein expressed in a heterologous system drastically decreased its pH sensitivity.^9^ Based on this finding, we have produced mice with the H165A mutation to determine whether loss of NCX1 pH regulation alters cardiac electrophysiology and intracellular Ca handling in mouse ventricular cardiomyocytes. We find that H165A mice are healthy, fertile and resistant to ischemia/reperfusion injury. However, they also exhibit significant changes in cardiac repolarization, ion channel function and Ca regulation. This is the first evidence that pH regulation of NCX1 affects cardiac physiology.

## Methods

### Data Availability

All data supporting this study are available from the corresponding author upon reasonable request.

The Institutional Animal Care and Use Committee at Cedars-Sinai Medical Center approved all animal protocols. Experimental group sizes using both male and female mice were based on previous studies of NCX1 knockout and overexpression in mice.^6, 17, 18^ Using CRISPR/Cas9, we generated H165A mice to render NCX1 resistant to pH and to assess the effects of this resistance on heart function, cellular Ca regulation and excitation-contraction (EC) coupling. Ventricular function was studied using 2D and M-mode echocardiography along with simultaneous surface electrocardiography (ECG). qPCR and western blotting were used to assess expression of key genes and proteins involved in Ca regulation, pH regulation and EC coupling. Enzymatically isolated adult ventricular myocytes were loaded with Ca and pH indicators and studied with the whole cell perforated or ruptured patch clamp techniques to measure action potentials, NCX1 and L-type Ca currents, Ca transients and SR Ca load. Statistical tests for experiments shown in each figure are reported in the corresponding legend. An expanded methods section is provided in the Supplemental Materials, as are uncut gel blots and detailed listing of PCR primers. Data are presented as mean ± standard deviation (SD).

## Results

### Generation and baseline characteristics of H165A mice

We used CRISPR/Cas9 to produce a global knock-in mutation of the NCX1 gene (Slc8a1) on a mixed C57Bl/6Hsd-C57Bl/6NCrl background, in which the mature protein has an alanine instead of histidine at position 165 (CAT to GCT). We also introduced a silent mutation in an upstream codon for arginine (AGG to AGA) to both simplify PCR differentiation between knock-in and wildtype NCX1 and to abolish the Cas9 PAM sequence. Successful mutation and germline transmission were verified by both PCR and DNA sequencing in founders and subsequent generations. Mice were then bred to homozygosity from proven mutants. We verified successful global knock-in of the codon for alanine corresponding to position 165 in the NCX1 protein (along with a silent mutation for arginine in the preceding codon as a convenient marker) in progeny of the CRISPR/Cas9 founders by sequencing tail genomic DNA. Next, we used qPCR to confirm the mutation was present in heart tissue from H165A mice. Relative to the housekeeping gene GAPDH, there was no expression of the wildtype codon for arginine (AGG) measured at the silent mutation site in H165A mutants (Figure 1B, *across silent mutation*), whereas RNA (GCT, corresponding to alanine at position 165) and protein expression (C2C12 Ab) of the mutant gene product remained at WT levels (Figure 1B, *downstream from silent mutation* and Figure 1C respectively).

**Figure 1.**
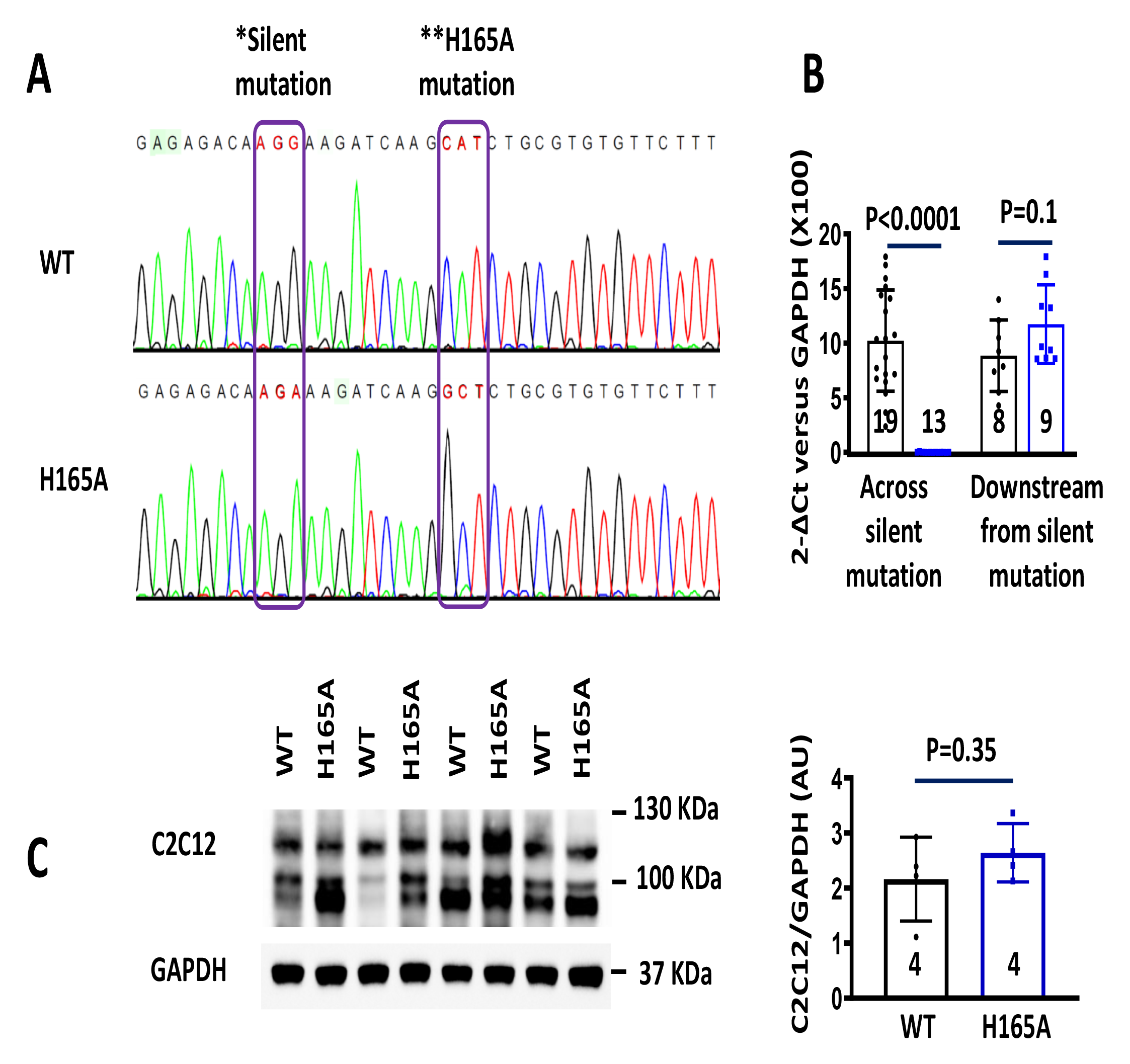
H165A pH-insensitive NCX mice produced by CRISPR/Cas9 technology. **A.** Genomic DNA sequencing of wildtype (WT) and mutant (H165A) NCX1 gene. Lefthand box outlines the upstream silent mutation site corresponding to arginine at position 161 (AGG to AGA). The righthand box indicates the histidine (CAT) to alanine (GCT) mutation at position 165. **B**. RT-qPCR verified that cDNA from H165A mice lacked the WT codon corresponding to arginine at position 161 (*across silent mutation*), but fully expressed the downstream codon corresponding to alanine at position 165. **C**. Western blots (*left*) using an antibody recognized by both WT and mutant NCX1 proteins (C2C12) revealed normal NCX1 expression levels as shown on the summary graph to the *right.* Summary data presented as Mean ± SD; unpaired t-test.

H165A mice live to adulthood and are fertile. There is no evidence of cardiac hypertrophy, with no change in heart weight to tibia length (HW/TL, Supplemental Figure 1). Unexpectedly, male H165A mice weigh less than age-matched WT (C57Bl6) mice (Figure 2A, *left*). Female H165A mice also weigh less than WT at 2.5 weeks, but this difference was no longer significant by 6 weeks of age (Figure 2A, *right*). Left ventricular (LV) systolic function by echocardiography under anesthesia is reduced modestly in H165A but still falls within normal limits (Ejection Fraction [EF] in WT: 64.6±9.1%, n=20; in H165A: 54.3±9.4%, n=13, p=0.004; Figure 2B). Surface ECG in anesthetized mice showed no change in heart rate or morphology, but QTc interval was decreased in H165A (from 25.0±6.9 ms in WT [n=12] to 19.8±3.5 ms in H165A [n=16], p=0.03, Figure 2C).

**Figure 2.**
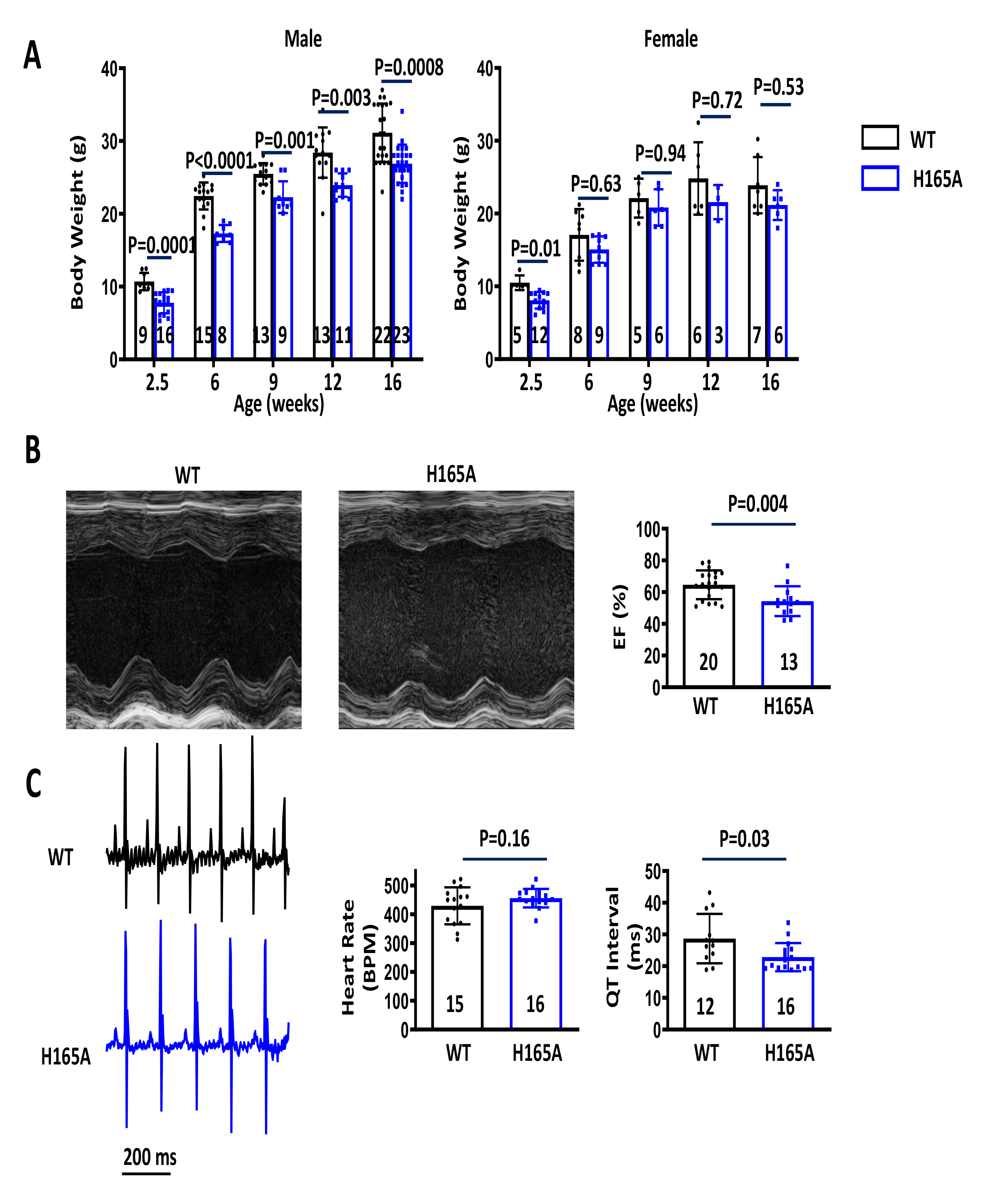
Body weight, echocardiography and electrocardiography. **A**. Mouse body weights from 2.5 to 16 weeks. Both male (*left*) and female (*right*) H165A mice (*blue*) exhibit delayed growth, but there is no statistical difference in female body weight by 6 weeks (Data presented as mean ± SD using two-way ANOVA with Sidak’s multiple comparison). **B**. M-mode echocardiography showing representative LV short axis views from WT and H165A mice. Bar graph to the *right* shows that LV function in H165A was modestly reduced, but still within normal range. **C**. Representative electrocardiograms from WT and H165A mice (*left*), and summary graphs showing no change in mean heart rate but a significant decrease in QT interval. (Data for B and C presented as mean ± SD with unpaired t-test).

To determine if there were changes in expression of common genes involved in excitation-contraction coupling, Ca handling, and acid/base regulation, we performed qPCR in isolated ventricular myocytes. Aside from a 32% reduction in Ca_V_1.2 gene expression, there were no significant changes in NBC, NHE, SERCA, RyR2, PMCA4, K_V_4.2, Na_V_1.5 or CaMKII (n=5-8 each group; Figure 3A; primers listed in Supplemental Figure 2). Protein expression profiles were determined by Western blots, which revealed increased expression of repolarizing channel proteins including K_V_2.1 and K_V_11.1 as well as increases in acid/base-regulating transporters NBC1 and NHE1, and the sarcolemmal membrane Ca transporter PMCA4; otherwise, there were no changes in expression of major ion channel and EC coupling proteins including Ca_v_1.2, Na_V_1.5, K_V_4.2, K_V_4.3, SERCA, PLB, RyR2, and CaMKII. (Figure 3B and Supplemental Figure 3).

**Figure 3.**
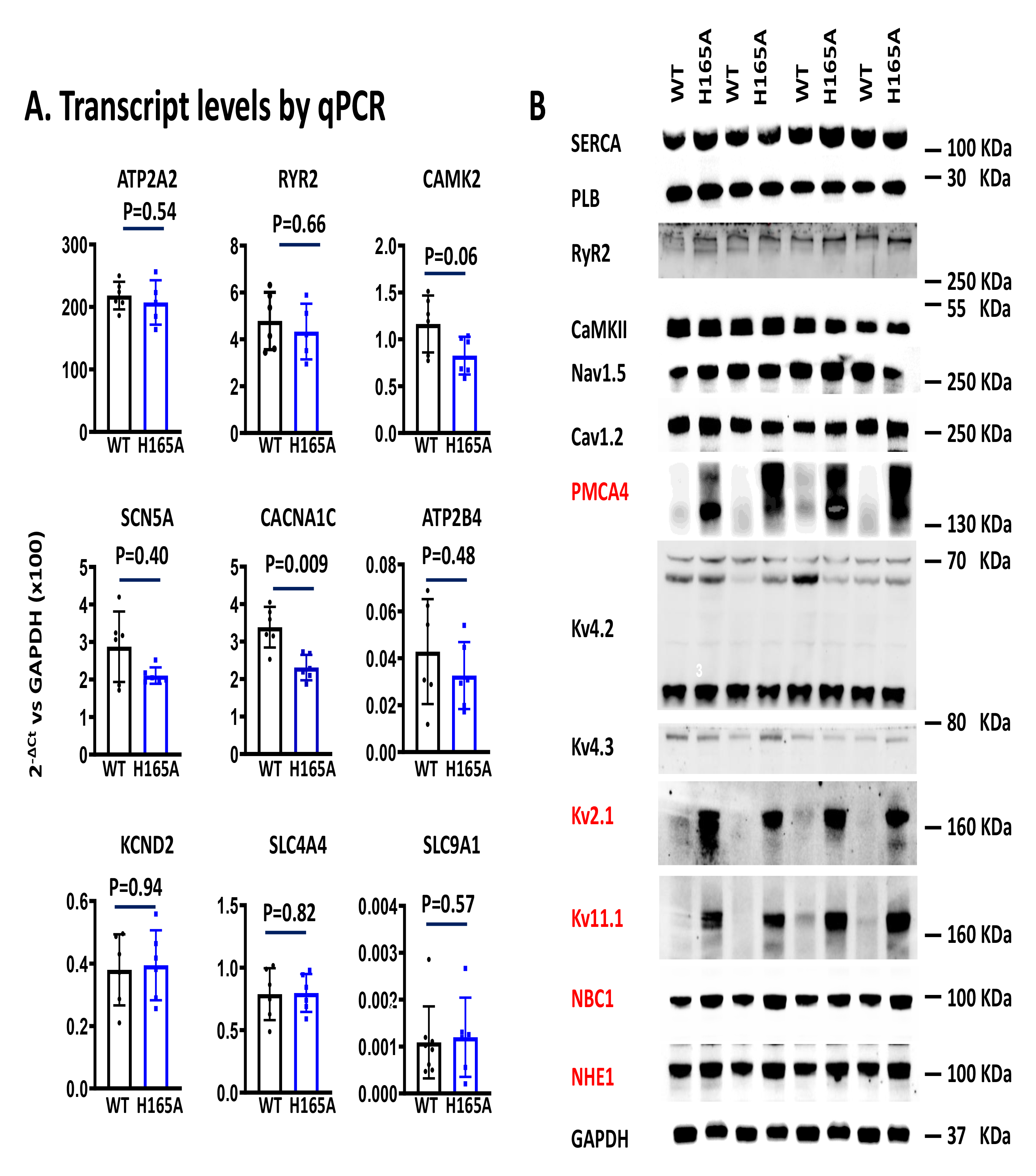
Expression of key excitation-contracton coupling and pH regulating genes and proteins. **A.** Summary graphs quantifying gene expression (Mean ± SD; Mann-Whitney test) measured by qPCR (2-ΔCt relative to the housekeeping gene GAPDH) in WT and H165A mice. Only CACNA1C, which codes for Ca_v_1.2 (L-type Ca channel), was reduced significantly in H165A. **B.** Protein expression by Western blots showing increases in expression of PMCA4, NBC1, NHE1 and several repolarizing K channel proteins (indicated by red font); otherwise there were no significant changes in H165A protein expression compared with WT. Protein quantification plots shown in supplemental Figure 3.

### Effect of H165A mutation on intracellular acidification

Given alterations in acid transporters, we examined the time course and extent of intracellular acidification in response to 80 mM Na-Acetate (Na-Ac) in intact cells. We perfused cells from WT and H165A mice with Tyrodes (1.8 mM Ca, pH 7.4) for one minute, then substituted 80 mM Na-Ac for equimolar NaCl for two minutes. When added to the bath, Na-Ac produces intracellular acidosis through influx of uncharged HAc across the plasma membrane followed by intracellular dissociation and an increase in free protons.^19, 20^ In response to Na-Ac, we found that intracellular pH decreased to the same extent (to ∼pH 6.5) in both WT and H165A myocytes in less than two minutes (n=29 cells from WT and n=33 from H165A; Supplemental Figure 4). Thus, we found no evidence that the H165A mutation altered the response of acid transporters to acidification, despite increased NBC1 and NHE1 protein expression.

### Electrophysiological properties of NCX Current in H165A mice and response to low pH

We assessed NCX current (I_NCX_) in patch clamped WT and H165A ventricular cardiomyocytes at pH 7.4. We applied voltage ramps from +80 mV to −120 (ramp speed: 680 mV/s) and recorded membrane current every 6 seconds for 1 minute before and after applying 5 mM Ni, as described previously (Figure 4A).^21^ After subtraction (Figure 4B), Ni-sensitive current (I_NCX_) was 35% smaller in H165A than WT at +80 mV (WT: 1.67±0.6 pA/pF; H165A 1.09±0.7 pA/pF [p=0.016], Figure 4C, *left*) and 40% smaller in H165A than WT at −120 mV (WT: −0.54±0.34 pA/pF; H165A: −0.32±0.3 pA/pF [p=0.009], Figure 4C, *middle*, n=36 in WT and n=28 in H165A). There was no difference in cell capacitance (Figure 4C, *right*).

**Figure 4:**
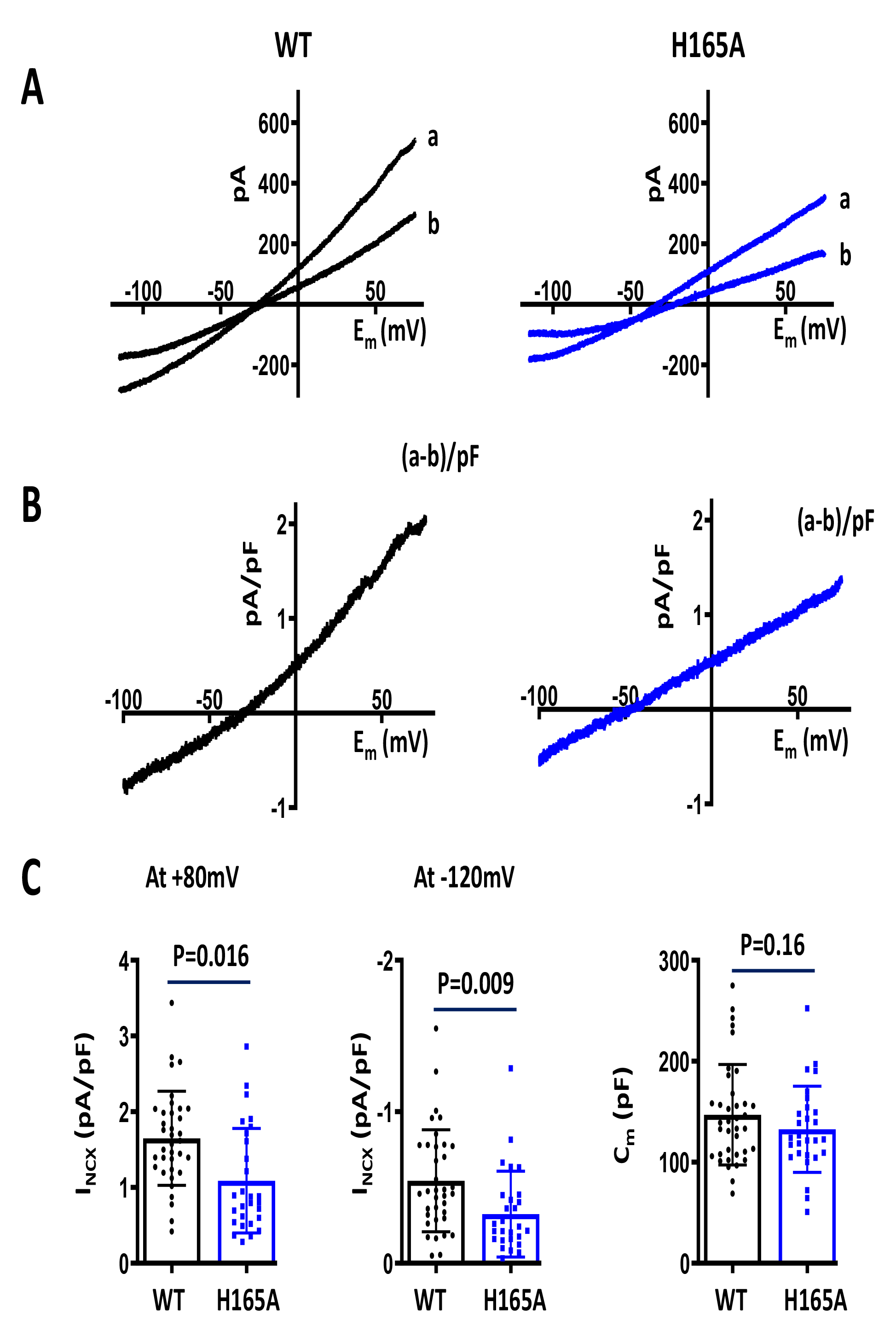
NCX1 current recordings in WT and H165A mice. **A.** Raw currents from WT and H165A myocytes recorded during voltage ramps from +80 mV to −120 mV before (a) and after Ni (b). **B.** Currents after Ni subtraction and normalization to cell capacitance (a-b)/pF. **C**. Summary plots showing reduced current density (pA/pF) at +80 mV and −-120 mV in the H165A mouse, but no change in cell capacitance (pF). Data presented as mean ± SD; nested t-test.

Next, we assessed the effect of acidosis on I_NCX_ by applying 80 mM Na-Ac for two minutes and then recording current in response to voltage ramps from +80 mV to −120 mV before and after adding Ni (Figure 5). Acidification with Na-Ac decreased I_NCX_ current at +80 mV by 69% in WT cells (n=6 from 5 mice, p=0.001, Figure 5B, *left*), but *had no effect* on I_NCX_ in H165A cells (n=9 from 5 mice, p=0.99). Similar results were noted at −120 mV (Figure 5B, *right*). These results establish that I_NCX_ in H165A ventricular myocytes is unaffected by acidification to ∼pH 6.5 by Na-Ac.

**Figure 5:**
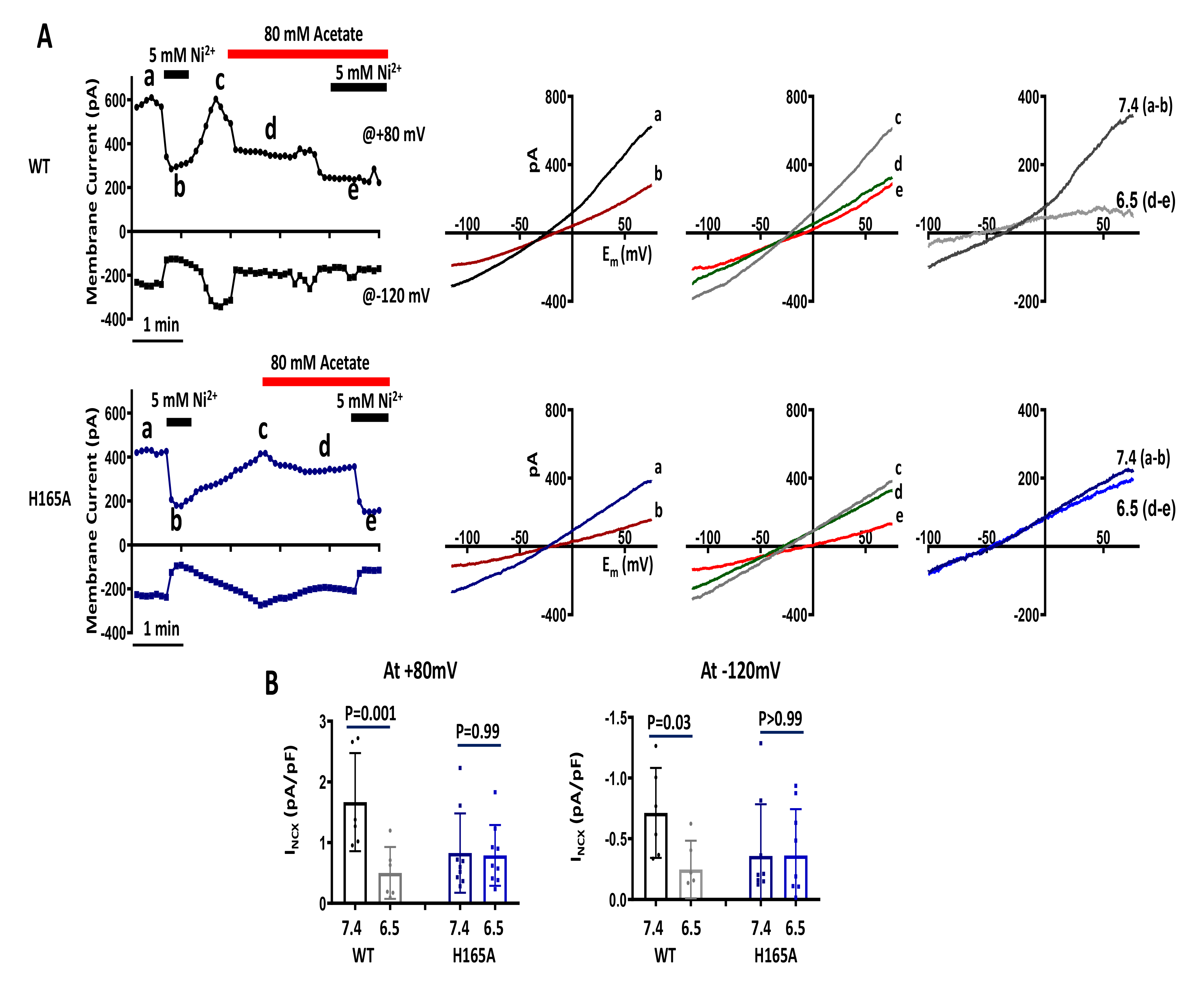
Effect of acidosis on NCX1 current in WT and H165A mouse. **A.** (*left*) Membrane current recorded at +80 mV and −120 mV during successive voltage ramps in representative WT and H165A mouse ventricular cardiomyocytes at pH 7.4 before (a) and after (b) applying 5 mM Ni^2+^ to block I_NCX_, then washing out Ni with pH 7.4 control bath solution (c) and applying 80 mM Na-Acetate bath solution to produce intracellular acidosis to pH 6.5 (d), and finally applying 5 mM Ni again to block I_NCX_ (e). Corresponding current-voltage plots for each step are shown to the *right.* Ni-subtracted currents (I_NCX_) are shown in the *far right* panel. **B.** Summary plots showing effect of acidification on NCX current at +80 mV and −120 mV (Mean ± SD; Two-way ANOVA with Tukey’s multiple comparison). Note that only WT current is suppressed by acidification.

### Calcium transients in H165A mice

To determine if EC coupling was altered by the H165A mutation, we measured intracellular Ca transients in ventricular myocytes loaded with fura-2 AM and field-stimulated by a MyoPacer (IonOptix) at 30 volts. When paced at 1 Hz, there was no significant difference in Ca transients between WT and H165A, including amplitude, systolic or diastolic Ca (Figure 6A, 6B). In separate experiments, we assessed sarcoplasmic reticulum (SR) Ca load by applying a 2 sec puff of caffeine (5 mM) to fluo-3 loaded patch-clamped myocytes (held at −80 mV) using a rapid solution exchanger.^18^ Just prior to applying caffeine, we equilibrated cells for 90 s using a series of conditioning pulses (200 ms clamps from −80 to 0 mV). In H165A cells, we found a 45% increase in the amplitude of the caffeine-evoked transient, consistent with elevated SR Ca load (Figure 6C).

**Figure 6:**
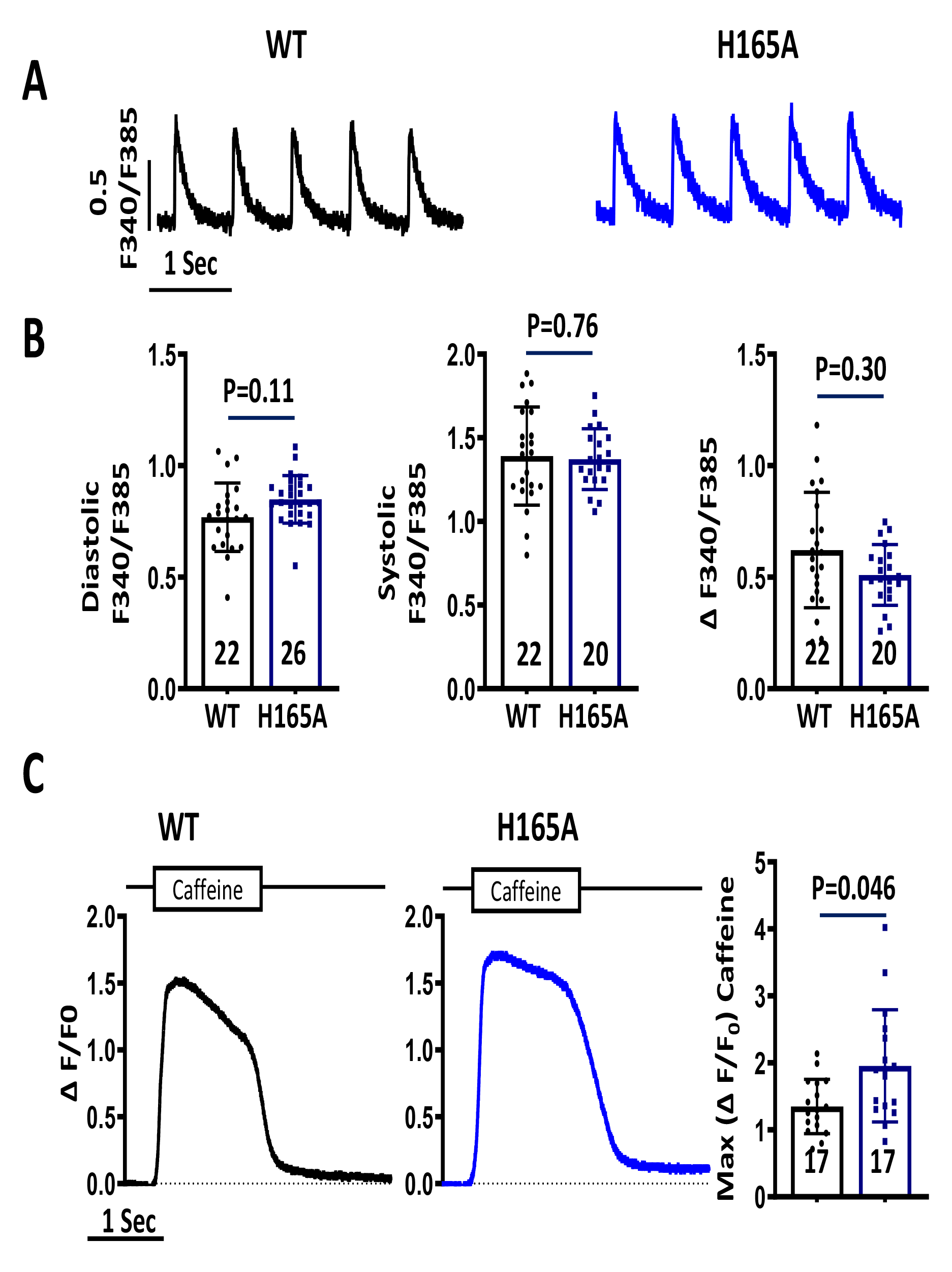
Ca transients in H165A mice. **A.** Fura-2AM fluorescence ratio traces recorded in representative ventricular myocytes from WT (*black*) and H165A (*blue*) mice during field-stimulation (1 Hz). **B.** Summary plots of diastolic, systolic and Ca transient amplitude (ΔF340/F385) during active pacing. **C.** Representative fluo-3 transients from WT (*left*) and H165A mice (*middle*) during rapid perfusion of 5 mM caffeine. Summary graph (right) showing higher fluo-3 fluorescence in response to caffeine in H165A cells, consistent with increased SR Ca load. Summary data presented as mean ± SD; nested t-test.

### Action Potential Characteristics

To further investigate the shorter QT interval in H165A, we recorded action potentials (APs) in isolated ventricular myocytes using the whole cell ruptured patch technique under current clamp mode. We found no difference in resting potential between WT and H165A. However, APD90 was markedly reduced in H165A mice (from 112.0 ± 76.6ms in WT [n=18/4 cells/animals] to 36.0 ± 25.2 ms in H165A [n=21/4 cells/animals], P=0.04) during pacing at 1 Hz (Figure 7A). There were no other changes in AP morphology. These findings are consistent with the reduced QT interval and increased expression of repolarizing K channels.

**Figure 7.**
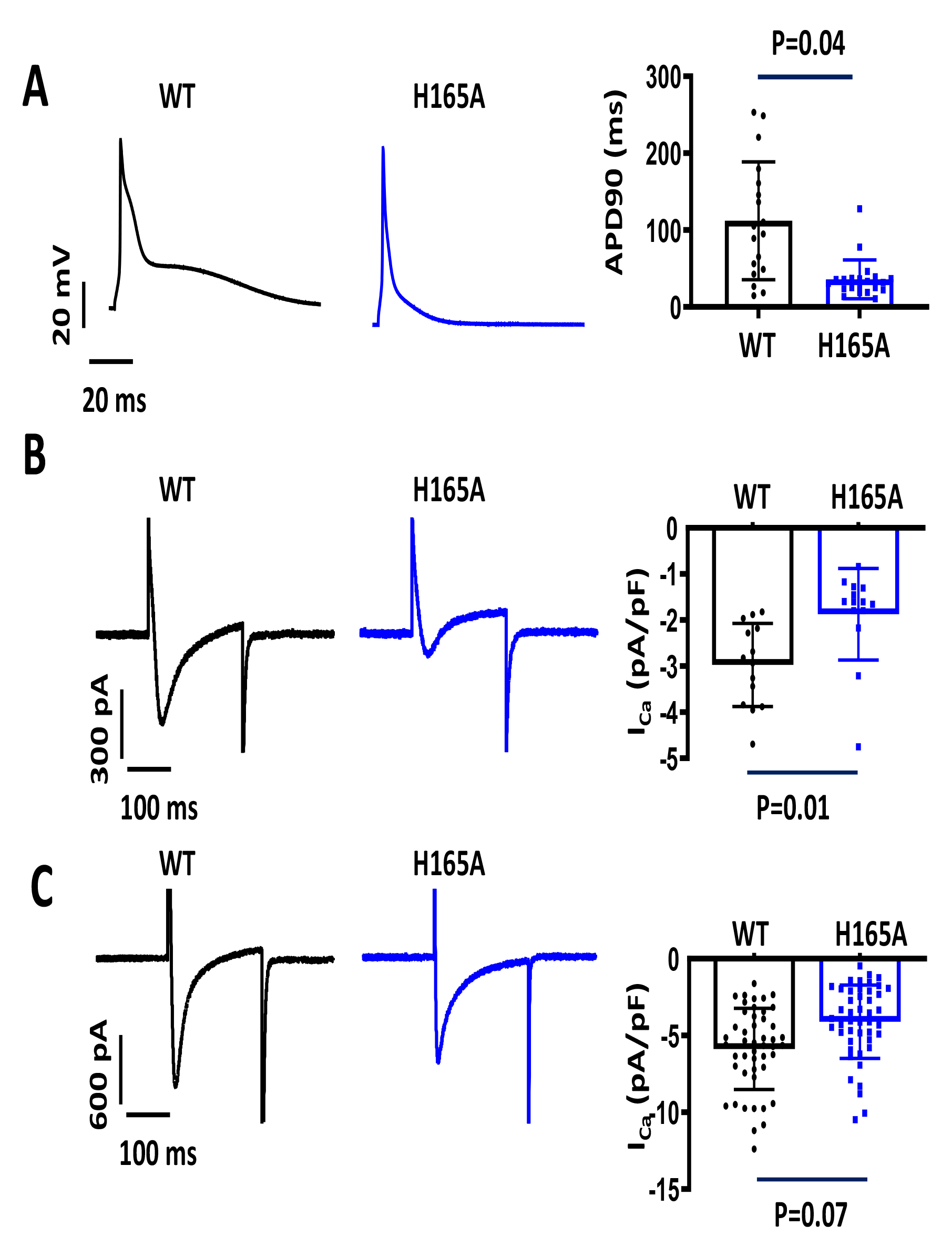
Action potentials and I_Ca_ in WT vs H165A. **A.** Representative action potentials (APs) from WT (*black*) and H165A (*blue*) patch-clamped ventricular myocytes. Note the reduction in AP duration (APD), which is summarized in the graph to the *right* (mean ± SD; Nested t-test) **B.** Representative traces of I_Ca_ recorded in WT and H165A myocytes during depolarization from −40 to 0 mV with either **B**, no Ca buffer in the pipette or **C**, 10 mM BAPTA included in the pipette. Summary graphs are shown to the *right* (mean ± SD; nested t-test).

### L-type Ca Current

To determine potential effects of the H165A mutation on the L-type Ca current (I_Ca_), we used the perforated patch technique to measure I_Ca_ in the absence of Ca buffering in patch clamped ventricular myocytes from WT and H165A mice. The external solution was modified Tyrodes with 1.8 mM Ca, but KCl was replaced by CsCl (5.4 mM). The high Cs internal solution is detailed in the Supplemental Methods. Cells were depolarized for 200 ms from a holding potential of −40 to 0 mV. I_Ca_ amplitude was reduced by ∼37% in the H165A mutant compared with WT (WT: −3.0±0.9 pA/pF, n=14/7 cells/animals; H165A: −1.9±1.0 pA/pF, n=14/7 cells/animals, p=0.01, Figure 7B). This is similar to the reduced I_Ca_ we observed previously in NCX1 KO cells, which is caused by elevated subsarcolemmal Ca.^22^ To test for this same mechanism in H165A mice, we recorded I_Ca_ in the ruptured patch configuration (in the absence of amphotericin) with 10 mM BAPTA included in the internal solution to buffer intracellular Ca. BAPTA eliminated the statistically significant difference in current amplitude between WT and H165A (WT: - 5.9±2.6 pA/pF, n=47/6 cells/animals; H165A: −4.1±2.4 pA/pF, n=46/6 cells/animals, p=0.07, Figure 7C), consistent with the hypothesis that elevated subsarcolemmal Ca further reduces I_Ca_ through Ca-dependent inactivation of L-type Ca channels.

### Ischemia/reperfusion studies

Ventricular-specific NCX1 KO mice are resistant to ischemia/reperfusion (I/R)^7^, and it has been hypothesized that pH regulation of NCX1 is an important component leading to I/R injury. We performed whole heart I/R studies in explanted hearts perfused in Langendorff fashion with oxygenated Tyrodes solution. Hearts were perfused with modified Tyrodes containing 1.8 mM Ca. 20 min of stop flow global ischemia was followed by 120 min reperfusion. Hearts were then removed, fixed in formaldehyde, sectioned, and stained with a 1% solution of 2,3,5-triphenyltetrazolium chloride (TTC). The area of necrosis was quantified using ImageJ to identify red (live) vs white (necrotic) tissue.^6^ H165A mice had a 67% reduction in the area of necrosis compared to WT (WT necrotic area: 25.9 ± 9.4% n=11; H165A area: 8.5 ± 3.8% n=7, p=0.0004; Figure 8A).

**Figure 8.**
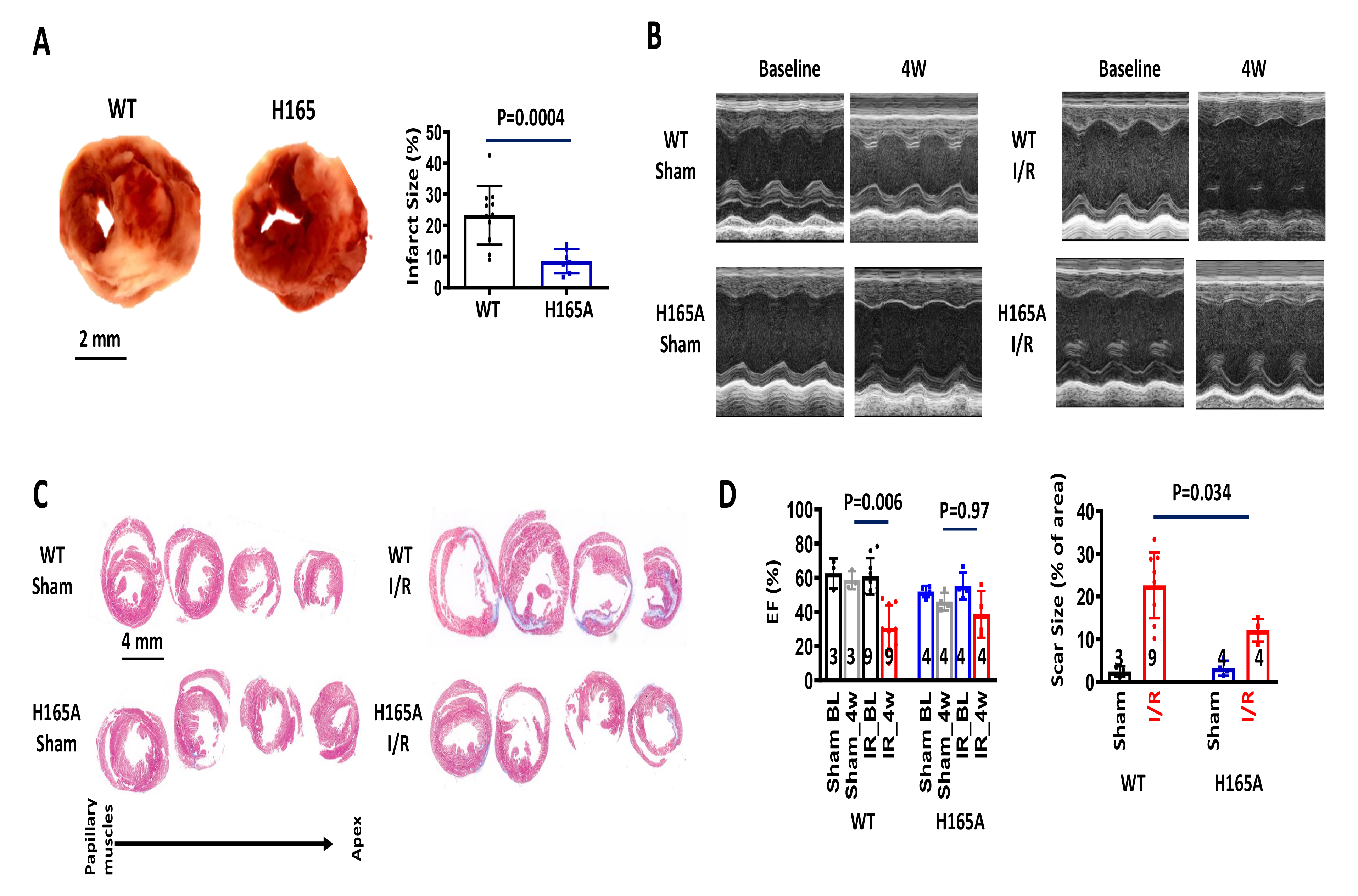
Ischemia/Reperfusion in H165A mice. **A.** Mouse heart sections from Langendorff-perfused hearts subjected to 20 min of stop flow ischemia followed by 120 min reperfusion. Sections stained with a 1% solution of 2,3,5-triphenyltetrazolium chloride (TTC) to quantify necrosis area. Summary graph (*right*) shows 67% reduction in infarct size in H165A compared to WT (mean ± SD; unpaired t-test). **B.** Representative M-mode echocardiogram images (short axis, LV) of WT and H165A mice before and 4W after 45 min ischemia followed by reperfusion produced by coronary artery ligation and release. **C.** The hearts were removed and fixed 4 weeks later, sectioned, and stained with Masson’s trichrome. **D.** Summary data of ejection fraction (EF) at baseline and 4 weeks after ischemia/reperfusion along with Sham controls (*left*). Quantification of the infarcted area as percentage of total heart tissue (*right*). There was far less fibrosis in H165A 4 weeks after ischemia/reperfusion than WT. Data presented as mean ± SD; three-way and two-way ANOVA.

We then performed coronary I/R *in vivo* using WT and H165A mice. 45 minutes of coronary occlusion were followed by reperfusion. Mice were allowed to recover for 4 weeks before undergoing echocardiography (Figure 8B) and histological assessment (Figure 8C). No mice died in either group. Compared to WT, H165A animals had significantly better preservation of left ventricular systolic function by echocardiography (Figure 8D, *left*). Furthermore, H165A hearts had a 46% reduction in scar size compared to WT (Figure 8D, *right*). Thus, the H165A mutation confers cardioprotection to I/R injury.

## Discussion

We used CRISPR/Cas9 to engineer a pH-insensitive NCX1 mouse to test the hypothesis that pH regulation of NCX1 is an essential component of cardiac physiology. H165A mice live to adulthood, breed normally, have no evidence of heart failure, and most importantly, I_NCX_ is unaffected by lowering pH (Figure 5). There are minimal changes in gene or protein expression, and Ca transients remain intact. Unexpectedly, we find that APD is substantially reduced, as is L-type Ca current, whereas SR Ca load is increased. Growth is reduced in males and delayed in females. When subjected to either global or regional coronary ischemia followed by reperfusion, pH-insensitive NCX1 mice have less I/R injury than WT.

Aside from the almost complete resistance of I_NCX_ to acidosis, the major electrophysiological consequence of the H165A mutation is the shortened APD and, consequently, reduced QT interval (Figures 2 and 7). Heart rate under light anesthesia is unchanged (Figure 2) and thus does not explain QT shortening. K_v_4.2, an important component of the transient outward current (I_TO_) and APD in mice ^23^ is also unchanged, but several other repolarizing K channels are increased (Figure 3). Although Ca_v_1.2 protein expression is normal (Figure 3), gene expression (Figure 3A) and I_Ca_ amplitude (Figure 7B) are both reduced, the latter in part a consequence of Ca-dependent inactivation. In this regard, we have observed reduced I_Ca_ in a different model, the ventricular-specific NCX1 KO mouse, despite normal protein expression.^5^ This is caused by increased subsarcolemmal Ca, a consequence of dramatically reduced Ca efflux in that model, resulting in Ca-dependent inactivation of L-type Ca channels.^22^ In H165A, the reduction in I_Ca_ is undoubtedly a major contributor to APD shortening. I_NCX_ amplitude is also reduced in H165A mice (Figure 4). While smaller I_NCX_ could further shorten the APD,^24^ it should be noted that some mouse AP models discount the importance of I_NCX_ on APD.^23, 25^

The mechanism underlying reduced I_NCX_ is not clear since NCX1^H165A^ protein expression is normal (Figure 1C). In a heterologous system (*Xenopus* oocytes), the H165A mutant elicits ionic currents with a magnitude comparable to those recorded from the WT exchanger, though in that model we can only effectively test transport in the “reverse” mode.^9^ These observations suggest that the H165A mutation may impact association with regulatory/partner proteins or trafficking of NCX1 to the membrane. The latter could be evaluated in the future using immunostaining.

EC coupling and intracellular Ca are dependent upon the balance of Ca influx and efflux mechanisms.^1, 26^ In H165A we have evidence for a new set point, specifically an increased SR Ca load, which helps preserve the Ca transient despite reduced trigger Ca entering through L-type Ca channels. This compensatory arrangement is distinct from the ventricular-specific NCX1 knockout mouse, where SR load is unchanged despite almost total elimination of NCX-mediated Ca efflux. One major difference in these models is the extent of compensation by Ca entry and efflux pathways, which is much more dramatic in the NCX1 knockout than it is in H165A.^27^

We have no explanation for the smaller size of H165A animals, but the finding suggests pH regulation of NCX1 is an important factor in growth and development. We speculate that low pH conditions during early embryogenesis inhibit NCX activity, thereby optimizing intracellular Na and Ca concentrations at critical milestones in development. Removing this control point with the H165A mutation disrupts Na and Ca balance, causing reduced heart and body weight in juvenile animals. Notably, the ventricular-specific NCX1 knockout mouse does not exhibit any developmental abnormalities. Thus, the phenotype is not linked to decreased expression of the exchanger. The sex difference in weight recovery suggests a hormonal component, which bears investigation in future studies.

### Ischemia/Reperfusion

NCX1 is an electrogenic transporter with a 3:1 stoichiometry (Na:Ca), and thus sensitive to altered concentrations of internal and external Na and Ca, as well as membrane voltage.^28, 29^ Under normal conditions, where intracellular Na is low and extracellular Na is high, NCX functions as the dominant Ca efflux mechanism (and Na influx mechanism) of cardiac myocytes.^30, 31^ Increased intracellular Na reduces Ca efflux by NCX (for example in response to cardiac glycosides), and at the extreme can reverse NCX to cause Ca influx^32^ and cellular injury.^7, 16, 33^ During ischemia, a dramatic increase in Na is mediated largely by two major acid-coupled Na influx transporters (NHE and NBC) in concert with reduced Na efflux by an energy deprived Na/K ATPase.^15, 34^ The low pH of ischemia inhibits NCX1 which prevents it from removing the excess Na *until* reperfusion abruptly restores pH *and* subsequently NCX1 function. Since the Na is so high, reverse NCX produces rapid and massive Ca influx. NCX1 *inhibition* to prevent Ca influx upon reperfusion has been proposed. Indeed several blockers of NCX reduced reperfusion injury in experimental models, though no pharmacological blocker of NCX has proven to be completely specific.^35–38^ We have shown that cardiac-specific NCX1 KO mice constructed with two different cre recombinase promotors (MLC2V and MerCreMer βMHC), are resistant to I/R injury.^6, 7^ However, NCX1 KO mice have reduce lifespans suggesting negative long-term consequences to its inhibition.^6^ Here we have shown that simply removing pH inhibition of NCX1, with its attendant compensations, is also effective at reducing I/R injury (Figure 8). We presume that maintaining NCX1 function throughout the I/R period avoids the excessive gain in Na during ischemia and the abrupt NCX reversal with rapid and overwhelming Ca entry upon reperfusion. The shortened APD and smaller I_Ca_, in concert with increased PMCA4 also help mitigate Ca loading under these conditions. Confirmation of this mechanism will require further experimentation.

In conclusion, the H165A mouse is the first evidence that pH regulation of NCX1 affects cardiac physiology and is a potential model for studying the role of NCX1 pH-sensitivity on both physiological and pathophysiological cardiac function, including cardioprotection. Growth retardation remains unexplained but our findings suggest NCX1 pH regulation is also important for development.

### Novelty and Significance

#### What is Known

- Sodium-calcium exchange (via NCX1) is the major calcium removal mechanism in cardiac myocytes, which is critical for regulating cellular calcium, contractility and rhythm.
- NCX1 is highly regulated by pH, a property implicated in the pathogenesis of ischemia/reperfusion injury.
- How pH regulation of NCX1 contributes to normal cardiomyocyte physiology is unknown.

#### What new information does this article contribute?

- We used CRISPR/Cas9 to engineer a novel pH-insensitive NCX1 mouse, by mutating Histidine to Alanine at position 165 of the NCX1 protein.
- NCX1 function is reduced 35-45% in these mutant mice, which are otherwise fertile and healthy with only modest reduction in LV function, shorter QT interval and sex-specific growth retardation. The mice are also resistant to ischemia/reperfusion.
- Ca transients are normal in H165A despite smaller I_Ca_, increased PMCA4, increased SR Ca load and shorter APD.
- This is the first evidence that pH regulation of NCX1 affects cardiac physiology. The H165A mouse can be used to study the role of NCX1 pH-sensitivity on both physiological and pathophysiological cardiac function.

## Non-standard Abbreviations and Acronyms

NCX1: Sodium-Calcium Exchanger isoform 1
Ca: Calcium
Na: Sodium
K: Potassium
ECC: Excitation-Contraction Coupling
RyR: Ryanodine Receptor
SERCA2: Sarcoplasmic Reticulum Ca2+ATPase 2
SR: Sarcoplasmic Reticulum
I/R: Ischemia/Reperfusion
GAPDH: Glyceraldehyde 3-phosphate dehydrogenase
EF: Ejection Fraction
PMCA4: Plasma membrane calcium ATPase 4
Na-Ac: Sodium-acetate
NBC1: Sodium bicarbonate cotransporter isoform 1
NHE1: Sodium hydrogen exchanger 1
PLB: Phospholamban

## References

1. Bridge JHB, Smolley JR and Spitzer KW. The relationship between charge movements associated with ICa and INa-Ca in cardiac myocytes. Science. 1990;248:376–378.

2. Reuter H. The Na+-Ca2+ Exchanger Is Essential for the Action of Cardiac Glycosides. Circulation Research. 2002;90:305–308.

3. Reuter H, Henderson SA, Han T, Ross RS, Goldhaber JI and Philipson KD. The Na+-Ca2+ exchanger is essential for the action of cardiac glycosides. Circ Res. 2002;90:305–8.

4. Pott C, Henderson SA, Goldhaber JI and Philipson KD. Na+/Ca2+ exchanger knockout mice: plasticity of cardiac excitation-contraction coupling. Ann N Y Acad Sci. 2007;1099:270–5.

5. Henderson SA, Goldhaber JI, So JM, Han T, Motter C, Ngo A, Chantawansri C, Ritter MR, Friedlander M, Nicoll DA, Frank JS, Jordan MC, Roos KP, Ross RS and Philipson KD. Functional adult myocardium in the absence of Na^+^-Ca^2+^ exchange: cardiac-specific knockout of NCX1. Circ Res. 2004;95:604–11.

6. Lotteau S, Zhang R, Hazan A, Grabar C, Gonzalez D, Aynaszyan S, Philipson KD, Ottolia M and Goldhaber JI. Acute Genetic Ablation of Cardiac Sodium/Calcium Exchange in Adult Mice: Implications for Cardiomyocyte Calcium Regulation, Cardioprotection, and Arrhythmia. J Am Heart Assoc. 2021;10:e019273.

7. Imahashi K, Pott C, Goldhaber JI, Steenbergen C, Philipson KD and Murphy E. Cardiac-specific ablation of the Na^+^/Ca^2+^ exchanger confers protection against ischemia/reperfusion injury. Circ Res. 2005;97:916–921.

8. Philipson KD, Bersohn MM and Nishimoto AY. Effects of pH on Na+ -Ca2+ exchange in canine cardiac sarcolemmal vesicles. Circ Res. 1982;50:287–293.

9. John S, Kim B, Olcese R, Goldhaber JI and Ottolia M. Molecular determinants of pH regulation in the cardiac Na+-Ca2+ exchanger. Journal of General Physiology. 2018;150:245–257.

10. Doering AE and Lederer WJ. The Mechanism by Which Cytoplasmic Protons Inhibit the Sodium-Calcium Exchanger in Guinea-Pig Heart-Cells. J Physiol-London. 1993;466:481–499.

11. Scranton K, John S, Escobar A, Goldhaber JI and Ottolia M. Modulation of the cardiac Na(+)- Ca(2+) exchanger by cytoplasmic protons: Molecular mechanisms and physiological implications. Cell Calcium. 2020;87:102140.

12. Boyman L, Hagen BM, Giladi M, Hiller R, Lederer WJ and Khananshvili D. Proton-sensing Ca2+ binding domains regulate the cardiac Na+/Ca2+ exchanger. J Biol Chem. 2011;286:28811–20.

13. Murphy E and Eisner DA. Regulation of intracellular and mitochondrial sodium in health and disease. Circulation Research. 2009;104:292–303.

14. Garciarena CD, Youm JB, Swietach P and Vaughan-Jones RD. H(+)-activated Na(+) influx in the ventricular myocyte couples Ca(2)(+)-signalling to intracellular pH. J Mol Cell Cardiol. 2013;61:51–9.

15. Tani M and Neely JR. Role of intracellular Na^+^ in Ca^2+^ overload and depressed recovery of ventricular function of reperfused ischemic rat hearts. Possible involvement of H^+^-Na^+^ and Na^+^-Ca^2+^ exchange. *CircRes*. 1989;65:1045-1056.

16. Imahashi K, Kusuoka H, Hashimoto K, Yoshioka J, Yamaguchi H and Nishimura T. Intracellular sodium accumulation during ischemia as the substrate for reperfusion injury. Circ Res. 1999;84:1401–6.

17. Torrente AG, Zhang R, Zaini A, Giani JF, Kang J, Lamp ST, Philipson KD and Goldhaber JI. Burst pacemaker activity of the sinoatrial node in sodium-calcium exchanger knockout mice. Proc Natl Acad Sci U S A. 2015;112:9769–74.

18. Kapoor N, Tran A, Kang J, Zhang R, Philipson KD and Goldhaber JI. Regulation of calcium clock-mediated pacemaking by inositol-1,4,5-trisphosphate receptors in mouse sinoatrial nodal cells. J Physiol. 2015;593:2649-63.

19. Swietach P, Leem CH, Spitzer KW and Vaughan-Jones RD. Experimental generation and computational modeling of intracellular pH gradients in cardiac myocytes. Biophys J. 2005;88:3018–37.

20. Spitzer KW, Ershler PR, Skolnick RL and Vaughan-Jones RD. Generation of intracellular pH gradients in single cardiac myocytes with a microperfusion system. Am J Physiol Heart Circ Physiol. 2000;278:H1371–82.

21. Lin X, Jo H, Sakakibara Y, Tambara K, Kim B, Komeda M and Matsuoka S. Beta-adrenergic stimulation does not activate Na+/Ca2+ exchange current in guinea pig, mouse, and rat ventricular myocytes. Am J Physiol Cell Physiol. 2006;290:C601–8.

22. Pott C, Yip M, Goldhaber JI and Philipson KD. Regulation of cardiac L-type Ca2+ current in Na+- Ca2+ exchanger knockout mice: functional coupling of the Ca2+ channel and the Na+-Ca2+ exchanger. Biophys J. 2007;92:1431–7.

23. Pott C, Ren X, Tran DX, Yang MJ, Henderson S, Jordan MC, Roos KP, Garfinkel A, Philipson KD and Goldhaber JI. Mechanism of shortened action potential duration in Na+-Ca2+ exchanger knockout mice. Am J Physiol Cell Physiol. 2007;292:C968–73.

24. Ferreiro M, Petrosky AD and Escobar AL. Intracellular Ca2+ release underlies the development of phase 2 in mouse ventricular action potentials. Am J Physiol Heart Circ Physiol. 2012;302:H1160–72.

25. Bondarenko VE, Szigeti GP, Bett GC, Kim SJ and Rasmusson RL. Computer model of action potential of mouse ventricular myocytes. American journal of physiology Heart and circulatory physiology. 2004;287:H1378–403.

26. Eisner DA. Ups and downs of calcium in the heart. J Physiol. 2018;596:19–30.

27. Pott C, Philipson KD and Goldhaber JI. Excitation-contraction coupling in Na+-Ca2+ exchanger knockout mice: reduced transsarcolemmal Ca2+ flux. Circ Res. 2005;97:1288–95.

28. Ottolia M, John S, Hazan A and Goldhaber JI. The Cardiac Na(+) -Ca(2+) Exchanger: From Structure to Function. Compr Physiol. 2021;12:2681–2717.

29. Hinata M, Yamamura H, Li L, Watanabe Y, Watano T, Imaizumi Y and Kimura J. Stoichiometry of Na+-Ca2+ exchange is 3:1 in guinea-pig ventricular myocytes. J Physiol. 2002;545:453–61.

30. Reuter H, Pott C, Goldhaber JI, Henderson SA, Philipson KD and Schwinger RH. Na(+)--Ca2+ exchange in the regulation of cardiac excitation-contraction coupling. Cardiovasc Res. 2005;67:198–207.

31. Despa S and Bers DM. Na(+) transport in the normal and failing heart - remember the balance. J Mol Cell Cardiol. 2013;61:2–10.

32. Despa S, Islam MA, Weber CR, Pogwizd SM and Bers DM. Intracellular Na(+) concentration is elevated in heart failure but Na/K pump function is unchanged. Circulation. 2002;105:2543–8.

33. Pike MM, Kitakaze M and Marban E. ^23^Na-NMR measurements of intracellular sodium in intact perfused ferret hearts during ischemia and reperfusion. AmJPhysiol. 1990;259:H1767–H1773.

34. Ch’en FF, Dilworth E, Swietach P, Goddard RS and Vaughan-Jones RD. Temperature dependence of Na+-H+ exchange, Na+-HCO3- co-transport, intracellular buffering and intracellular pH in guinea-pig ventricular myocytes. J Physiol. 2003;552:715-26.

35. Haigney MCP, Miyata H, Lakatta EG, Stern MD and Silverman HS. Dependence of Hypoxic Cellular Calcium Loading on Na+-Ca- 2+ Exchange. CircRes. 1992;71:547–557.

36. Ladilov Y, Haffner S, Balser-Schafer C, Maxeiner H and Piper HM. Cardioprotective effects of KB- R7943: a novel inhibitor of the reverse mode of Na+/Ca2+ exchanger. The American journal of physiology. 1999;276:H1868–76.

37. Takahashi T, Takahashi K, Onishi M, Suzuki T, Tanaka Y, Ota T, Yoshida S, Nakaike S, Matsuda T and Baba A. Effects of SEA0400, a novel inhibitor of the Na+/Ca2+ exchanger, on myocardial stunning in anesthetized dogs. Eur J Pharmacol. 2004;505:163–8.

38. Inserte J, Garcia-Dorado D, Ruiz-Meana M, Padilla F, Barrabes JA, Pina P, Agullo L, Piper HM and Soler-Soler J. Effect of inhibition of Na(+)/Ca(2+) exchanger at the time of myocardial reperfusion on hypercontracture and cell death. Cardiovasc Res. 2002;55:739–48.

